# Contribution of the Late Positive Potential to Emotional Memory: A Joint Neuro-Cognitive Approach to Memory Prediction

**DOI:** 10.64898/2026.10.16.759479

**Authors:** Robin Hellerstedt, Jordan Gunn, Deborah Talmi

**Author notes:** **Corresponding author:** Robin Hellerstedt.

## Abstract

Emotional events are better remembered than neutral events. The emotional Context Maintenance and Retrieval model (eCMR) explains this effect by simulating preferential encoding of emotional items, implemented by a free parameter that scales the learning rate. Incorporating a neural signal measured during encoding could make the model’s account more specific, permitting a prediction not only of average increased recall but also which emotional items are likely to be recalled. We investigated whether the late positive potential (LPP) event-related potential (ERP) effect could provide this item-specific information in a secondary analysis of EEG recorded while 38 participants encoded negative and neutral pictures for an immediate free recall test (Zarubin et al. 2020). First, we established that the LPP effect was modulated by emotion in an early (400-1000 ms) and a late (1000-2000 ms) time window and this modulation was reliable in 73.7% and 50% of the participants respectively. Second, we investigated at what level the LPP was related to subsequent memory. Emotion-enhanced LPP did not covary with emotion-enhanced memory between participants or across lists within participants, but the early LPP specifically predicted memory for emotional items on a single-trial level. Third, we compared eCMR variants to test whether trial-level early LPP should scale learning for both categories or only for emotional items, and whether it could replace or complement a common learning-rate increase for all emotional items. The best-fitting model combined emotion-specific LPP scaling with the common learning-rate increase. The finding that early LPP predicted enhanced recall only for emotional items is difficult to reconcile with accounts on which the LPP reflects attention or stimulus significance more generally, which predict a relationship to memory for neutral items as well. These results pave the way to quantitative prediction of internal recall trajectories by individual participants without requiring behavioural output.

## Introduction

Retrieved-context theory (Howard & Kahana, 2002) describes the rules that govern the chances that an experience encoded at one time point will come to mind at a later time point. The theory is supported by extensive behavioural (Healey & Kahana, 2016; Lohnas et al., 2015; Sederberg et al., 2008) and neural (Folkerts et al., 2018; Kragel et al., 2021; Manning et al., 2011; Polyn et al., 2005) evidence (for a recent review, see Lohnas & Healey, 2021) and offers the best explanation we have to date for the information processing mechanism the brain implements to give rise to memory recall. According to this theory, items become associated with a gradually-drifting internal context during encoding. This association functions bidirectionally, such that retrieving an item reinstates its associated context, which then cues further recall. Retrieved-context theory includes temporal and nontemporal dimensions of context, such as pre-experimental semantic associations and so-called “source” features –shared encoding conditions, such as the modality of item presentation or the orienting task performed during encoding. Emotional Context-Maintenance and Retrieval (eCMR, Talmi et al., 2019) showed that the emotional dimension is best described as a “source” feature. These models explain why participants recall contiguously items that evoke the same emotions, presented in the same modality or close in time to each other, and those encoded through the same orienting task, but none of these mechanisms account for enhanced recall of emotional items, a robust empirical phenomenon (Dolcos et al. 2025; Kensinger & Ford, 2020; Talmi, 2013). This manuscript focuses on the mechanism that allows eCMR and following models (CMR3, Cohen & Kahana, 2022; TCM-SR, Zhou et al. 2025) to explain enhanced emotional memory - that emotional arousal should be simulated as a modulator of the learning rate.

It is known that emotional items are preferentially processed during encoding (Anderson, 2005; MacKay et al., 2004; Pourtois et al., 2013; Schupp et al., 2006). eCMR and subsequent models simulate this by scaling the learning rate for associations from context to item features during encoding, thereby simulating stronger binding between emotional items and their encoding context. The degree of extra processing is governed by the value of parameter *φ*_emot_., and governs whether emotional items are modelled as receiving extra processing during encoding. This mechanism gives rise to the robust finding that participants who encode a mix of emotional and neutral items recall the emotional items better (Dolcos et al. 2025; Kensinger & Ford, 2020;Talmi, 2013) and the less-intuitive finding that when emotional items are encoded and recalled separately from neutral items, their advantage is attenuated (Talmi, Lohnas & Daw, 2019).

Using an independent neural measure to model variation among items, rather than relying only on a free parameter shared by all emotional items, would provide a more specific account of emotional memory and a stronger test of the theory (Turner et al., 2016). An excellent candidate is the late positive potential (LPP), an event-related potential (ERP) effect that increases in amplitude when participants process emotionally-significant visual stimuli (Cuthbert et al., 2000; Hajcak et al., 2010; Schupp et al., 2000, 2006). Because eCMR attributes the emotional encoding advantage to preferential processing during study, and the LPP is sensitive to the enhanced processing of motivationally significant stimuli, it is natural to consider whether LPP variation can provide this item-level information. In this project, our aim was to examine the suitability of the LPP for neuro-cognitive modelling of memory recall. We pursued this aim in three steps. First, we establish the individual-level reliability of the LPP under memory-experiment conditions. Second, we test whether and how LPP variation predicts emotional memory. Finally, we investigate how best to incorporate the neural measure into the computational model.

The LPP is broadly characterised as reflecting affective stimulus significance (Hajcak & Foti, 2020; Olofsson et al., 2008), and its emotional modulation has been observed at the group level across a range of stimuli and presentation durations. However, for the LPP to serve as a useful candidate marker of emotional encoding, it must also differentiate emotional from neutral processing at the individual level. Recently, Schupp and Kirmse (2021) examined individual-level sensitivity to emotional content using a case-by-case approach across three studies with different emotional stimulus categories. They observed that the LPP was larger for high-compared to low-arousing stimuli in 98% of individual-level tests. A complementary specificity analysis confirmed that this effect was driven by emotional content rather than low-level stimulus differences. Additionally, the emotional modulation of the LPP showed good internal consistency (Moran et al., 2013) and was reliable within individuals across testing sessions (Weinberg et al., 2021).

For the LPP to serve as a useful neural model input, however, the same individual-level reliability must hold under conditions typical of memory experiments. Although emotional modulation of the LPP has been observed at the group level in intentional-encoding paradigms followed by free recall (Barnacle et al., 2018, Zarubin et al., 2020), it is not yet known whether comparable individual-level sensitivity is preserved. Participants in Schupp and Kirmse (2021)’s study viewed briefly-presented pictures passively. By contrast, participants in typical memory recall experiments take part in multiple study-test cycles, and are therefore normally aware of an upcoming memory test. Intentional encoding instructions have important impact on memory-relevant operations (Popov & Dames, 2023), which could attenuate or obscure the emotional modulation of the LPP in several ways. Participants may shift attention from processing each stimulus in isolation to comparing stimuli or organising them for later recall, adopt encoding strategies that reduce processing differences between emotional and neutral items, or engage in rehearsal that redistributes attention more evenly across stimulus types. The potential impact of an encoding “task set” is supported by evidence that task demands modulate emotional ERP effects (Schindler & Straube, 2020), that encoding instructions alter the organisation of recall (Healey et al., 2019), and that attention manipulation can increase or eliminate the recall advantage of emotional items (Talmi et al., 2007; Talmi & McGarry, 2012). Our first aim, therefore, was to replicate Schupp and Kirmse (2021)’s findings using a setup more typical in memory research. In addition to testing whether the LPP’s emotional modulation survives intentional encoding, a replication would increase confidence in the generality of the findings. Relative to Schupp and Kirmse (2021), the present study uses a different stimulus set, a longer presentation duration (2000 ms vs. 150 ms), and a larger sample (*n* = 38 vs *n* = 16-18 per experiment).

Attention and memory are typically coupled, such that when all else is held equal, increased attention during encoding increases participants’ ability to recall attended stimuli subsequently (Chun & Turk-Browne, 2007). Increased attention to emotional stimuli during encoding is considered a key driver of enhanced memory for these stimuli (Mather & Sutherland, 2011; Talmi, 2013). The LPP has been interpreted as a signal of increased stimulus processing as a consequence of stimulus significance (Hajcak & Foti, 2020) or tagging of memories as important for consolidation (Fields, 2023). Accordingly, previous studies report that emotional arousal increased both the LPP and memory (Dolcos & Cabeza, 2002, Zarubin et al., 2020). Yet direct tests of whether the LPP–memory relationship is consistent within or between participants are scarce (see Fields, 2023 for a review of the literature on the link between the LPP and memory). Indeed, Barnacle et al. (2018) found evidence attention-related LPP modulation and downstream memory advantage can be functionally dissociated: the emotional modulation of the LPP was equally strong in pure and mixed lists under intentional encoding conditions, while emotional memory enhancement depended on list composition. Understanding how LPP variation relates to memory within a mechanistic model may help bridge this gap.

The relationship between LPP at encoding and subsequent memory could depend on the level of analysis. If we can establish that the LPP is reliable enough at the individual-level, the second step would, therefore, be to differentiate participant-level, list-level and item-level relationship between the LPP and subsequent memory. At the between-participant level, individuals with larger emotion-dependent LPP increases may exhibit larger emotion-dependent memory enhancement. At the within-participant level, emotion may enhance memory most for study lists where encoding emotional items was accompanied by a larger LPP. A dissociation between within- and between-participant effects in ERPs has been demonstrated for the left parietal old-new effect, where robust within-participant effects were observed despite null between-participant effects (MacLeod & Donaldson, 2017). Clearly, if trial-level information underpins the relationship, then averaging across participants or lists could also decrease ability to observe a relationship between the LPP and emotion-enhanced memory, so an analysis at the level of the single trial may be necessary. Practically, however, the LPP may be too noisy at the single-trial level to be useful for modelling purposes. Establishing whether the relation between the LPP and emotion-enhanced memory is better captured at the single-trial level, when averaged over items in a list, or over the entire session is necessary to add the LPP to eCMR, and is therefore central to the present manuscript.

If the second step is successful, and we establish not only that the LPP is a reliable signal but also the level at which its relationship to memory holds, the final step is to investigate how best to deploy this signal as a model input, to improve the ability of the model to predict memory recall. We therefore compared variants of eCMR in which preferential learning of emotional items and the LPP could make separate contributions to encoding strength. The model comparison assessed whether including the LPP improved prediction of which items were recalled, whether LPP-based modulation substituted for or complemented category-wide preferential learning of all emotional items, and whether the LPP–recall relationship was shared across emotional and neutral items or specific to emotional items.

### Empirical investigation

We report a secondary data analysis of data from Zarubin et al. (2020). We re-analysed data from their mixed list condition (Figure 1), where participants encoded neutral items and negative, arousing items (referred to as ‘emotional items’ below). After encoding each list, participants were asked to describe the items they could recall to the experimenter. Zarubin and colleagues reported that emotional items were recalled more frequently than neutral ones. We classified items by their subsequent memory status as recalled or forgotten, with a resulting experimental design crossing the factors Emotion and Subsequent memory. A more detailed description of the methods in the original experiment is provided in the original paper.

**Figure 1.**
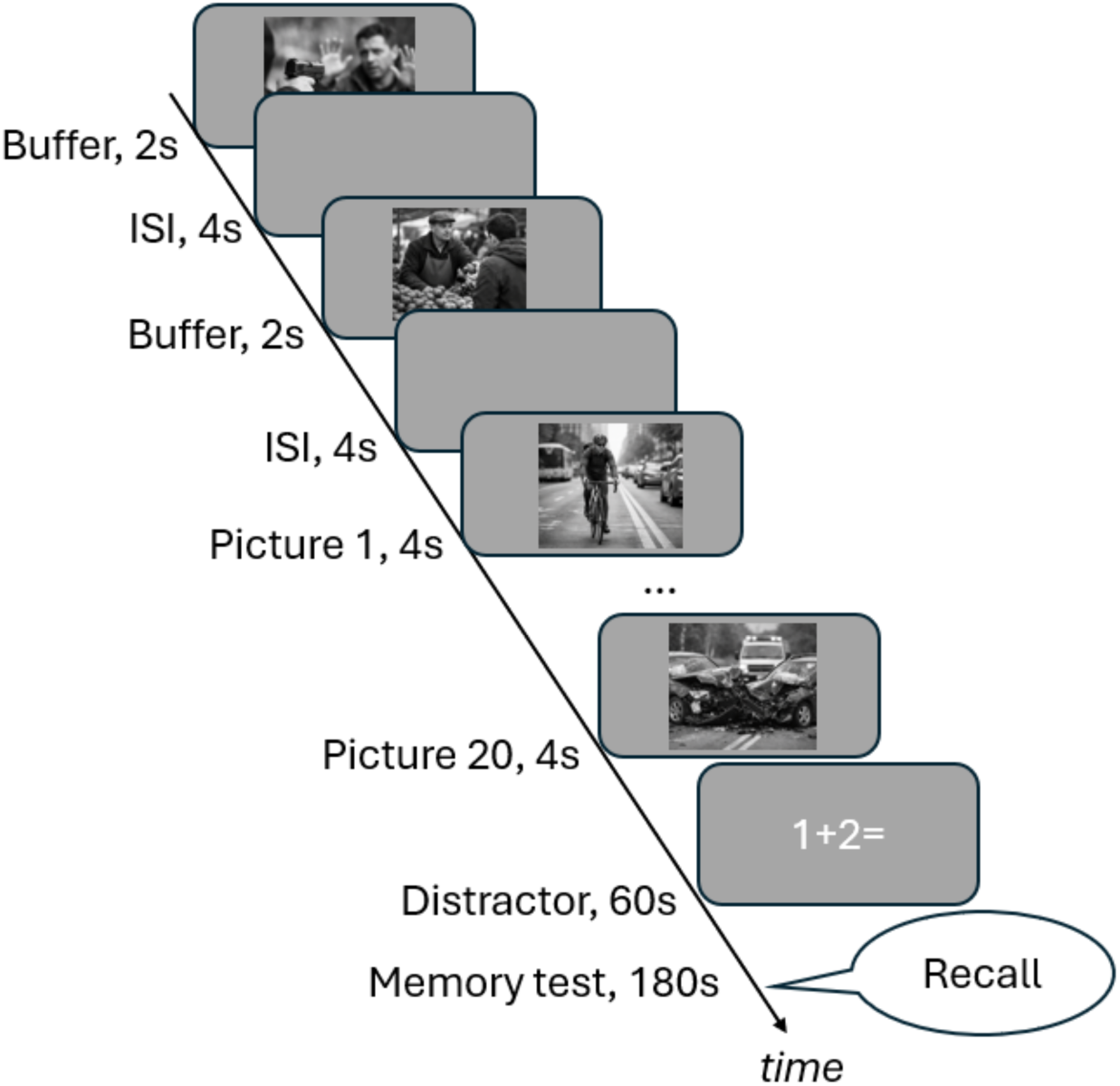
Timeline of list encoding and the memory test in the mixed-list condition of Zarubin et al. (2020). Each list comprised 22 pictures, and there were nine lists in total. The first two pictures in each list served as buffer items and were excluded from all behavioural and EEG analyses. The images shown here were generated with ChatGPT 4 and are provided for illustration only.

## Method

### Participants

We analysed data from the forty participants included in the mixed condition in Zarubin et al. (2020). We excluded one participant who did not have data from three lists and one participant who had fewer than 16 trials in one of the conditions after EEG preprocessing. The final sample consisted of 38 participants (28 females, *M*age = 20, *SD*age = 1.07). Gender and age data was missing for one of the participants. All participants provided written informed consent and the experiment was approved by Wofford College Institutional Review Board (Zarubin et al., 2020).

### Materials

There were 22 pictures in each list and a total of nine lists. The first two pictures were used as fillers to control for primacy effects and were excluded from data analysis. The pictures were selected from the International Affective Picture System (Lang et al., 2005), the Geneva Affective Picture Database (Dan-Glauser & Scherer, 2011), the Emotional Picture Set (Wessa et al., 2010), the image pool of Talmi et al. (2007) and Google Images. The pictures were either of negative valence or neutral valence. The neutral condition was divided into semantically unrelated and semantically related neutral pictures in the original study, but they were collapsed in this secondary data analysis to increase the number of trials in the neutral condition.

### Procedure

The participants studied 22 pictures shown consecutively on the screen for 2000 ms with a 4000 ms inter-trial interval, intended to reduce emotional carryover effects (Talmi & McGarry, 2012). The presentation order of both the lists and the images within lists was randomised. A one-minute arithmetic distracter task followed immediately after the study task to reduce rehearsal of the pictures and recency effects. A verbal recall test followed the distracter task. The participants were instructed to recall images only from the list that they had just seen. The participants had three minutes to recall the images from the study phase but could stop sooner if they had no more items to report.

### EEG pre-processing

Raw EEG files were downloaded from the open science framework page for the Zarubin et al. (2020) study. The EEG was pre-processed with the EEGLAB toolbox in MATLAB (Delorme & Makeig, 2004) using a standard semi-automated pipeline (for a similar preprocessing pipeline see Hellerstedt et al. 2023). The data was re-referenced offline to the average of the left and right mastoid electrodes and filtered between 0.1 and 40Hz with a band pass FIR filter. The data was time-locked to the stimulus and segmented into epochs from −2000 ms to 3000 ms post stimulus onset. The epochs were baseline corrected using a pre-stimulus time window from −200 ms to 0 ms. Customized threshold functions from the FASTER toolbox (Nolan et al., 2010) were used to detect and reject bad channels and bad epochs based on participant-level z-transformed values over trials and electrodes that exceeded ± 3. Independent component analysis was used to correct for EOG artefacts, by manually removing components classified as blinks, horizontal or vertical eye-movements. Bad channels which were deleted prior to ICA were interpolated after the ICA cleaning. In addition, bad channels within single epochs were detected and interpolated with a criterion of ±3 SD. All participants contributed at least 16 trials in each of the four ERP conditions. The mean number of trials was 30.7 (*SD* = 6.08) in the negative remembered condition, 26.1 (*SD* = 5.54) in the negative forgotten condition, 48 (*SD* = 15.33) in the neutral remembered condition and 66.5 (*SD* = 16.93) in the neutral forgotten condition after pre-processing. Trials that were rejected during EEG preprocessing were excluded from the behavioural analyses as well to ensure that the EEG and behavioural data contained the same trials and to increase the chance of finding a relation between the two measures.

### Data Analysis

#### Behavioural analysis

We investigated whether there was an emotion enhanced memory effect by comparing the proportion of negative items recalled and the proportion of neutral items recalled with a paired *t*-test. Zarubin et al. (2020) included two neutral conditions (semantically related and semantically unrelated pictures), so this analysis confirmed that the emotion-enhanced memory effect was still present once these conditions were collapsed.

#### Focal ERP Analysis

The purpose of this analysis was to replicate the LPP ERP effect. We analysed the electrodes and time windows where the LPP is typically maximal in the literature. Averaged EEG amplitude was extracted from four centroparietal electrodes, Cz, CP1, CP2 and Pz, in an early time window between 400–1000 ms and a late time window between 1000–2000 ms post stimulus onset. Below we refer to these data as early and late LPP amplitude, respectively. We extracted the EEG amplitudes separately for subsequently remembered and subsequently forgotten trials for both the negative and the neutral condition. EEG amplitudes were z-transformed against the mean and standard deviation of the −200 to 0 ms pre-stimulus interval. Standardisation was performed at the single-trial level, separately for each electrode. The z-transformed LPP was then extracted for both the early and the late time window from the same centro-parietal electrodes used in the focal ERP analysis. We refer to these data below as the standardised LPP amplitudes. All mixed-effects model analyses were conducted using the lme4 package in R (Bates et al., 2015).

We first performed a linear mixed-effects model to analyse if LPP amplitude was predicted by emotion and subsequent memory. In this model we used LPP amplitude as the dependent variable, Emotion (Negative vs Neutral) and Subsequent memory (Remembered vs Forgotten) as fixed effects, and random intercept + random slope effects of participant. The formula was:

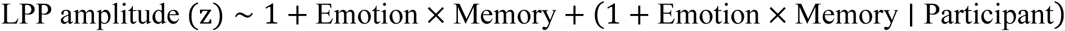

The random effect structure needed to be simplified to only include a random intercept effect of participant in the late LPP time window due to a warning of singular fit. The statistical significance of the fixed effects and their interaction was assessed using *χ*^2^ Wald test as implemented in the car package (Fox & Weisberg, 2019). Significant interactions were followed-up using pairwise contrasts as implemented in the emmeans package in R (Lenth, 2024) applying a Benjamini & Hochberg (false discovery rate, FDR) correction for multiple comparisons.

#### Global ERP Analysis

We complemented the focal analysis with a global, data-driven analysis of all data points between 0 and 2000 ms, the interval during which the stimulus was presented. This analysis served two purposes. First, it tested whether the LPP effect was also present under a more conservative procedure, one that makes no a priori assumptions about the electrode sites or latencies at which the effect should occur. Second, because the focal analysis was restricted to a small set of electrode sites and latencies, it could not detect effects arising elsewhere in the spatiotemporal space; the global analysis was not subject to this constraint and could detect if other ERP effects are more relevant for predicting emotion and memory. In this analysis we controlled for multiple comparisons using cluster-based permutation tests as implemented in the FieldTrip toolbox (Maris & Oostenveld, 2007; Oostenveld et al., 2011). The first step of this analysis is to perform *t*-tests at every ERP data sample. Significant samples, uncorrected at *α* = .05 (two-tailed), that were neighbours in time or space, including at least two electrodes, were grouped into clusters. The cluster level t-value was computed by calculating the sum of the t-values from all samples included in the cluster. Finally, we compared the cluster level *t*-value in the observed data to the size of the maximal cluster in 5000 permutations, for which the datapoints were randomly swapped between conditions within participants. The cluster level Monte-Carlo *p*-value was calculated as the proportion of the permutations for which the maximum cluster was larger than the clusters in the observed data. The exact spatio-temporal distribution of the clusters should be interpreted with caution given that the correction for multiple comparisons are applied at the cluster level rather than at the individual sample level (Sassenhagen & Draschkow, 2019).

#### Bootstrap Analysis

Having established that the emotional modulation of both the early and the late LPP was significant in the focal and global ERP analyses, we next used a bootstrap analysis to assess, separately for each participant, the reliability of the negative–neutral difference in early and late LPP amplitude. The rationale of this analysis is that negative stimuli should elicit a more positive LPP than neutral stimuli independent of which specific trials contributed to calculating the ERPs. The number of trials was first equated for the two conditions with under-sampling. The data included in this analysis was extracted from the same centro-parietal electrode sites as in the focal ERP analysis, i.e. Cz, CP1, CP2 and Pz. Bootstrap resampling from single trials was used to create a distribution of 1000 resampled ERPs per condition. A sliding time window was applied to detect the most positive LPP peak (the mean amplitude of the 100 ms time window between 400–1000 ms for the early LPP and between 1000–2000 ms for the late LPP).

Difference scores were calculated by subtracting the amplitude in the neutral condition from the amplitude in the negative condition for each bootstrap sample. Next, the proportion of positive bootstrap differences were calculated for the two time-windows based on the rationale that participants with robust LPP effects should have a large proportion of positive bootstrapped differences. We interpreted the emotion enhanced LPP as being reliable if a proportion of 0.9 or more of the bootstrapped differences were positive because this cutoff has been used in similar ERP analyses in the concealed information test literature (for example Rosenfeld, 2020).

#### Between-participants associations

For the between-participants analysis of the relationship between the LPP and emotion-enhanced memory, we calculated six scores per participant: the average standardised early LPP amplitude, the average standardised late LPP amplitude, and the average memory performance across the nine lists, each computed separately for the emotional and the neutral condition. We then computed the difference scores, subtracting the average performance in the neutral condition from performance in the emotional condition, arriving at three difference scores per participant, for early LPP amplitude, late LPP amplitude, and memory performance, which were then related using Bayesian Pearson correlations.

#### Within-participants associations

For the within-participants analysis, we computed for each participant the Pearson correlation between emotion-enhanced LPP and emotion-enhanced memory across the nine lists, separately for early and late LPP. The resulting correlation coefficients were then tested against zero with a Bayesian one-sample *t*-test, again separately for early and late LPP.

#### Trial-level associations

We used a binomial generalized linear mixed-effects model to investigate if Subsequent memory was predicted by Emotion and LPP amplitude at the single trial level. In this model we used Subsequent memory as the dependent variable and included Emotion (Negative vs Neutral) and LPP amplitude as fixed effects and random intercept + random slope effects of Participant. Two models were conducted, separately for the early and late LPP time windows. In each case, the model formula was:

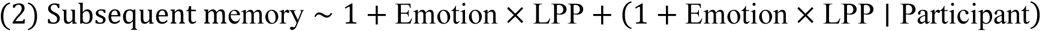

## Results

### Behavioural Results

The paired *t*-test showed that memory performance was significantly higher for negative items (*M* = 0.54, *SD* = 0.1) compared to neutral items (*M* = 0.42, *SD* = .14), *t*(37) = 6.18, *p* < .001.

### Focal ERP Analysis

We next conducted a focal ERP analysis to test whether the LPP at centro-parietal electrode sites differentiated negative from neutral items in the early and the late time window, and whether it was related to subsequent memory. We therefore fitted a linear mixed-effects model with Emotion (Negative vs. Neutral) and Subsequent memory (Remembered vs. Forgotten) as predictors of LPP amplitude, separately for each time window (Figure 2). In the early time window, there was a main effect of Emotion (***β*** = .20, ***χ*^2^**(1) = 38.01, *p* < .001) and a main effect of Subsequent memory (***β*** = .25, ***χ*^2^**(1) = 26.70, *p* < .001) and a significant Emotion × Subsequent memory interaction (***β*** = .10, ***χ*^2^**(1) = 14.70, *p* < .001). Follow-up pairwise contrasts showed that the LPP amplitude was higher for subsequently remembered than for subsequently forgotten negative items (*z* = −5.61, *p* < .001), while there was no difference in LPP amplitude for subsequently remembered and forgotten neutral items (*z* = −.88, *p* = .379).

**Figure 2.**
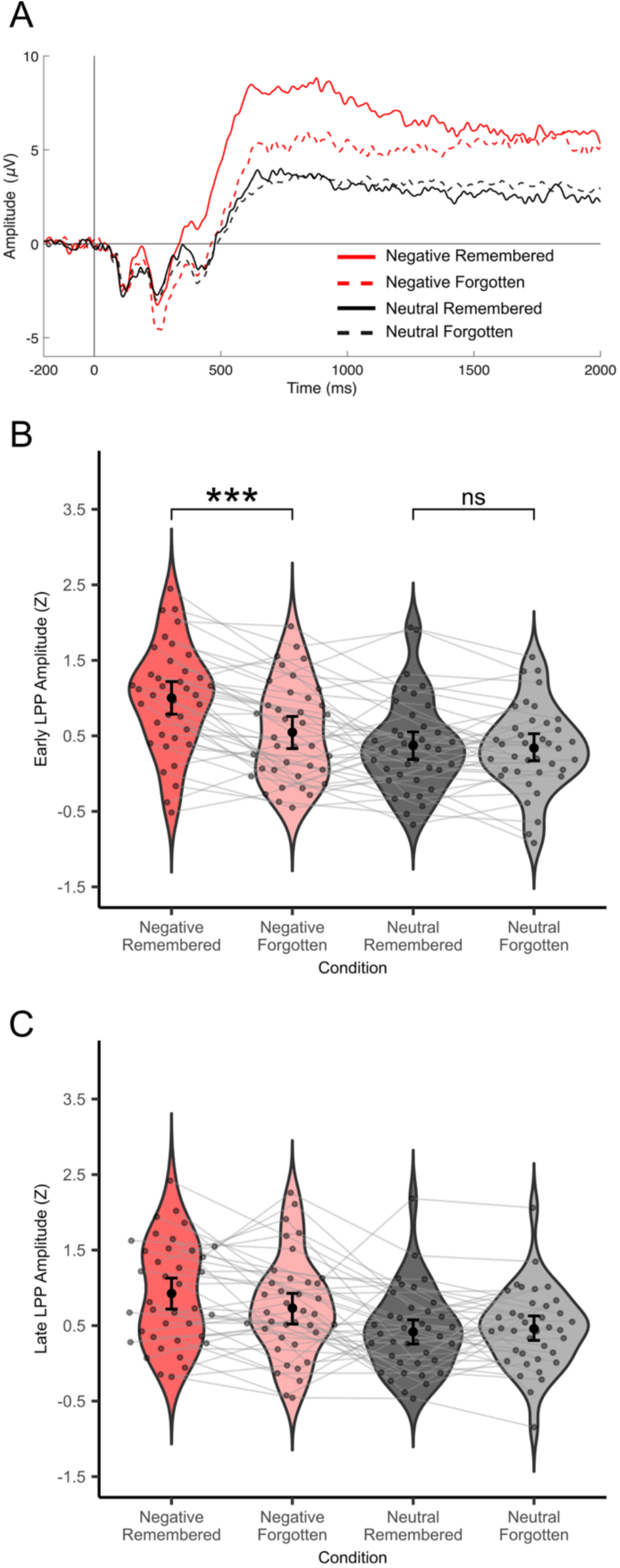
Focal ERP analysis. A) Event-related potentials (ERPs) from the encoding phase of the experiment averaged from the region of interest included in the focal ERP analysis (electrodes Cz, CP1, CP2, Pz). Conditions: red = negative items, black = neutral items; solid line = subsequently remembered items, dashed line = subsequently forgotten items. B) Z-transformed ERP amplitudes from the early LPP time window (400-1000 ms post stimulus presentation). Grey dots indicate means from individual participants, the black dot indicate the group mean and the error bar shows the 95% confidence interval of the mean. Light grey lines display what condition means comes from the same participant. Significance levels: *** < .001, * < .05. C) Z-transformed ERP amplitudes from the late LPP time window (1000-2000 ms post stimulus presentation). No follow-up contrasts were computed as the Emotion × Subsequent memory interaction was not significant in this time window.

In the late time window, there was a main effect of Emotion (***β*** = .18, ***χ*^2^**(1) = 46.39, *p* < .001), indicating that the LPP amplitude was higher for negative than for neutral items. There was no significant main effect of Subsequent memory (***β*** = .04, ***χ*^2^**(1) = 2.13, *p* = .144). There was a trend for a significant Emotion × Subsequent memory interaction (***β*** = .05, ***χ*^2^**(1) = 3.06, *p* = .080).

### Global ERP Analysis

We complemented the focal analysis with a global, data-driven analysis of all data points in the 0–2000 ms stimulus interval. Because it makes no a priori assumptions about electrode sites or latencies, this analysis provided both a more conservative test of the LPP effect and a means of detecting effects elsewhere in the spatiotemporal space. First, it tested whether the LPP effect was also present under a more conservative procedure, one that makes no a priori assumptions about the electrode sites or latencies at which the effect should occur. Second, the global analysis can detect effects outside of the regions and time windows of interest in the focal analysis in the spatiotemporal space. Results are displayed in Figure 3. We first analysed the main effect of Emotion (Negative vs Neutral trials) and found a positive cluster (*p* < .001) which was significant between approximately 380-2000ms with a widespread distribution. In other words, there was an emotionally enhanced LPP effect which roughly overlapped with the early and late time windows analysed in the focal analysis.

**Figure 3.**
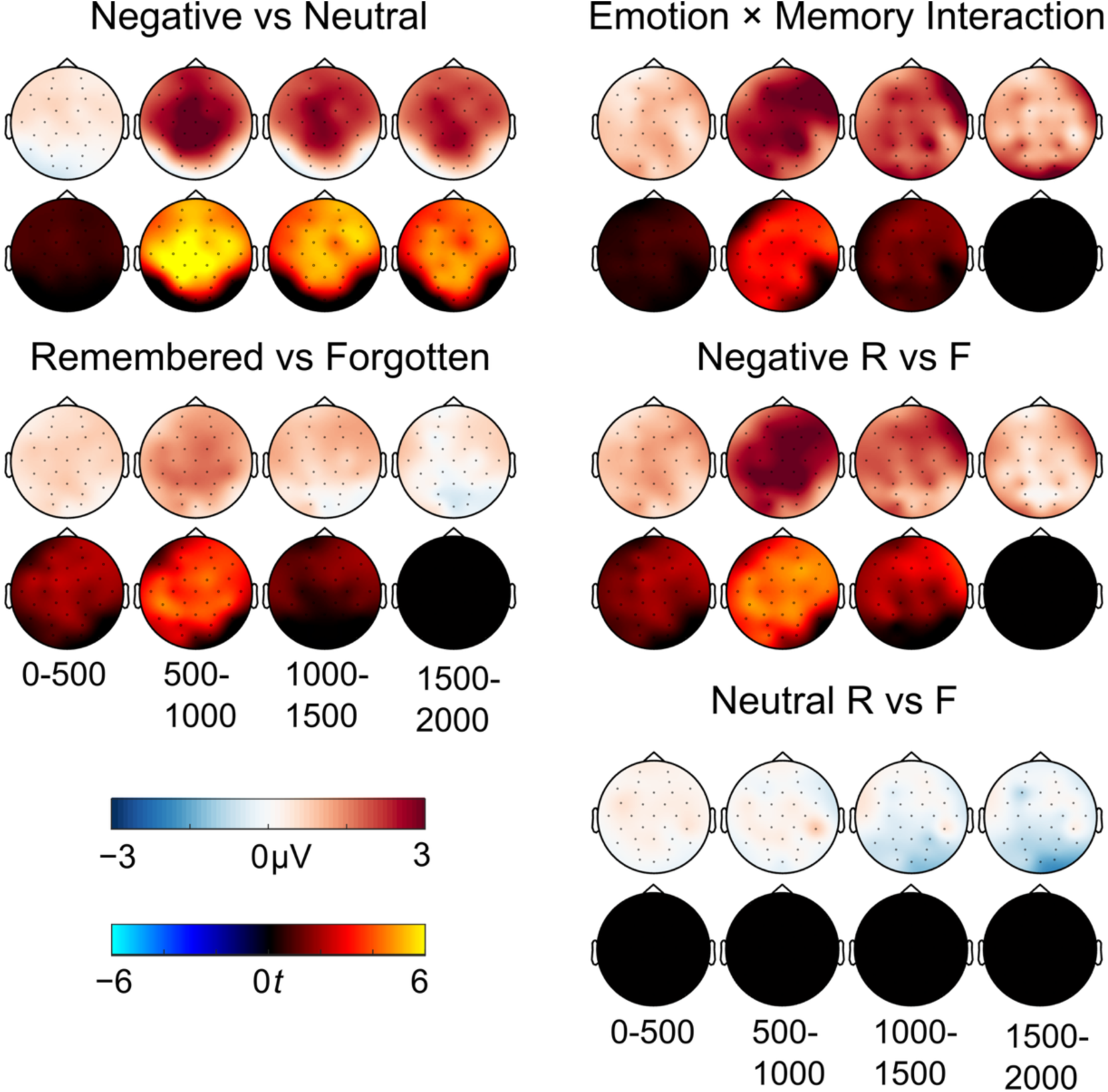
Global ERP analysis. The upper rows (blue to red colourmap) shows amplitude differences between conditions, in the encoding phase of the experiment, in micro volts in the 0-500 milliseconds (ms), 500-1000 ms, 1000-1500 ms and 1500-2000 ms time windows. Lower rows (cold to hot colourmap) show thresholded topographic maps of *t*-values from significant clusters from the global ERP analysis for the same time windows. Abbreviations: R = subsequently remembered, F = subsequently forgotten.

Replicating the focal analysis, there was also a main effect of subsequent memory with more positive ERPs for remembered compared to forgotten trials (*p* < .001) with a widespread spatial distribution between around 210-1240ms.

In addition, there was an Emotion × Subsequent Memory interaction between 360-1240ms (*p* < .001) with a widespread distribution, indicating that the negative items subsequent memory effect was larger than the neutral items subsequent memory effect. In addition, there was a widespread cluster (*p* = .029, two tailed alpha = .025) between 1240-1390ms that did not meet the significance threshold. Follow-up cluster permutation tests showed that for negative items, remembered items were related to more positive going ERPs between 220-1420ms (*p* < .001). This effect was also widespread. In contrast, there was no significant subsequent memory ERP effect for neutral items (all *p*s > .19).

Overall, the global analysis replicated the focal analysis and shows that these results hold up in more conservative tests with correction for multiple comparisons. The analysis also shows that the time windows and electrodes analysed in the focal analysis capture the maximum of the LPP effects well in this data set.

### Bootstrap Reliability of LPP amplitudes

Having established a significant LPP modulation of emotion in the focal and global ERP analyses we next performed a bootstrap analysis to test if this emotional LPP effect was reliable within participants. This analysis showed that the early LPP was reliable in 73.7% of the participants while the late LPP effect was reliable in 50% of the participants (Figure 4A).

**Figure 4.**
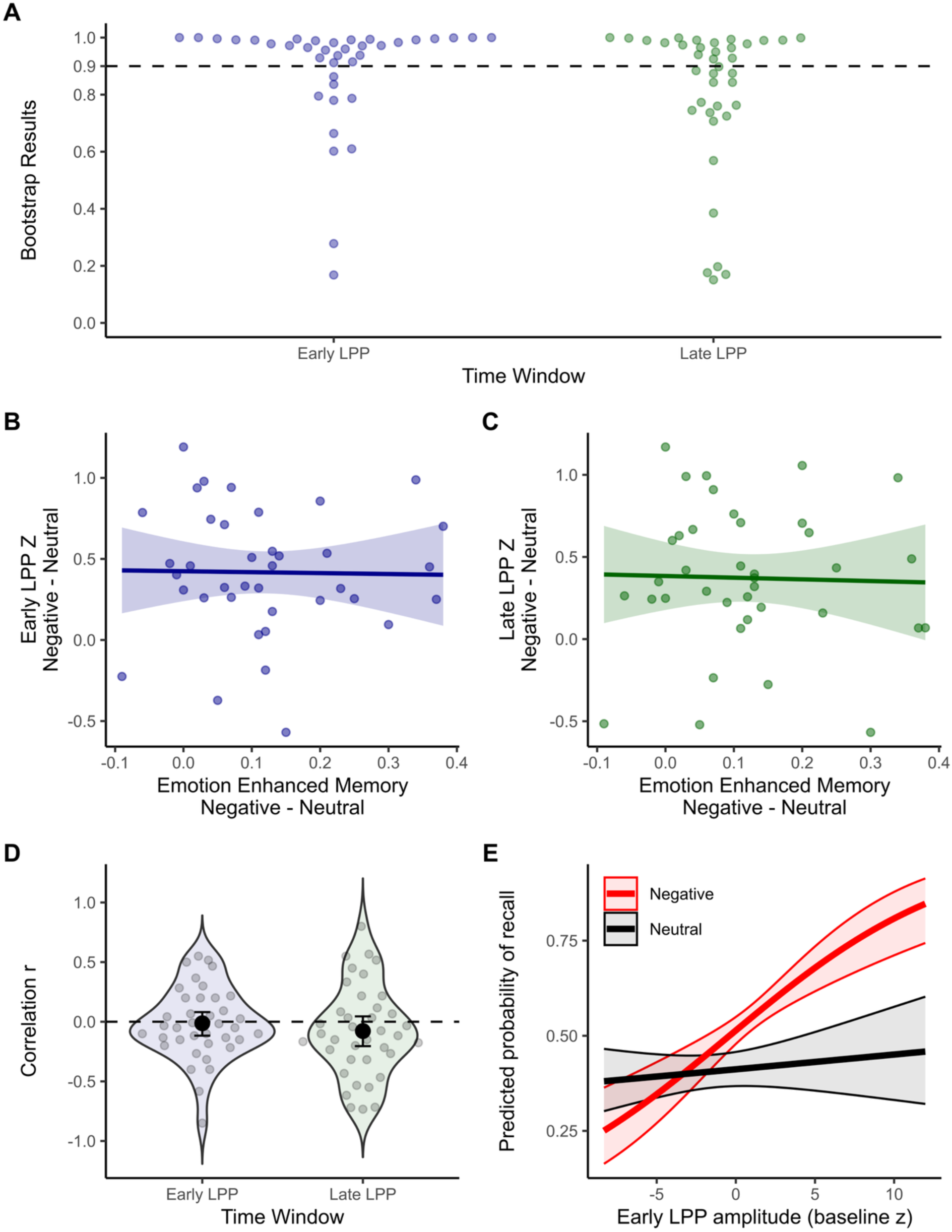
Late positive potential reliability and relationship to subsequent memory. A) Results from the bootstrap analysis depicting the proportion of bootstrap resampling iterations where the amplitude for the negative condition was more positive than for the neutral condition. The dashed line indicates the reliability threshold set at 0.9. Early LPP time window = 400 ms to 1000 ms, Late LPP time window = 1000 ms to 2000 ms. B) Between subjects Pearson correlation between the z-transformed amplitude difference between negative and neutral conditions and the emotion enhanced memory effect for the early LPP time window. C) The same for the late LPP time window. D) Within participant correlation between LPP amplitude and subsequent memory for the early LPP and the late LPP time windows. E) Early LPP amplitude predicts recall for negative but not neutral items. Fitted values from the binomial generalized linear mixed-effects model predicting subsequent memory from Emotion, Early LPP amplitude (z), and their interaction

#### Relationship between the LPP and Emotion Enhanced Memory: Between and within-participant associations

For the LPP to be useful as a model input, it needs to relate to memory performance at a relevant level of analysis. We therefore examined that relation at the between-participant, within-participant, and single-trial levels in turn. We first investigated whether the emotionally enhanced LPP amplitude (negative minus neutral items) correlated with emotionally enhanced memory (negative minus neutral items) between participants (Figures 4B and 4C). The Bayesian Pearson correlation showed moderate evidence for no correlation for both the early time window (*r* = –.018, *BF*01 = 4.925) and the late time window (*r* = –.028, *BF*01 = 4.885). Next, we investigated the correlation between emotion enhanced LPP amplitude and emotion enhanced memory within participants (Figure 4D). There was moderate support for no correlation in both the early time window (*t*(37) = –.26, *p* = .795, *BF*01 = 7.658) and the late time window (*t*(37) = – 1.26, *p* = .215, *BF*01 = 3.698).

#### Trial-level associations

Finally, we conducted binomial generalized mixed-effects models to test if Emotion and LPP amplitude predicted memory (Figure 4E). In the early time window, there was a main effect of Emotion (***β*** = .24, *χ*^2^(1) = 29.36, *p* < .001), a main effect of LPP (***β*** = .08, *χ*^2^(1) = 18.87, *p* < .001) and an Emotion × LPP interaction (***β*** = .06, *χ*^2^(1) = 11.20, *p* < .001). We unpacked the interaction by running the same analysis separately for negative and neutral items:

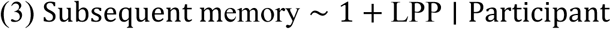

For negative items, there was a significant positive relationship between early LPP amplitude and memory (***β*** = .14, *χ*^2^(1) = 25.43, *p* < .001). In contrast, there was no significant relationship between early LPP amplitude and memory for neutral items (***β*** = .01, *χ*^2^(1) = .48, *p* = .488). In the late time window, there was a main effect of Emotion (***β*** = .23, *χ*^2^(1) = 66.58, *p* < .001), showing once again that memory was higher for negative than neutral items, but importantly there was neither a main effect of LPP amplitude (***β*** = .02, *χ*^2^(1) = 1.98, *p* = .160) nor any interaction between Emotion and LPP amplitude (***β*** = .02, *χ*^2^(1) = 2.44, *p* = .118).

## Discussion

In sum, the focal and the global ERP analyses converged in showing that emotion modulated the LPP in both time windows, and interacted with subsequent memory in the early time window only. The early LPP was reliable in a larger proportion of participants than the late LPP (73.7% versus 50%), and only the early LPP predicted recall at the single-trial level, selectively for negative items. Neither time window was associated with emotion-enhanced memory in aggregated scores. We therefore incorporated the LPP into eCMR at the single-trial level using standardised amplitudes from the early time window, matching the level and time window at which the LPP–memory relationship was observed.

### Computational modeling of the early LPP contribution to emotional recall

The behavioral and EEG analyses identified two forms of variation in recall. At the group level, emotional items were recalled more often than neutral items. At the trial level, Early LPP was more strongly associated with subsequent recall for emotional than neutral items. The empirical analyses distinguish these patterns statistically, but they do not determine how EEG voltages at this early time window should be mapped onto the encoding mechanism that produces emotional memory enhancement. We therefore tested whether using this signal to scale learning could replace the same increase in learning for every emotional item, or whether the two forms of modulation explained complementary variation in recall.

We organized this comparison around three focused questions. First, we tested whether allowing all emotional items to share the same adjustment to learning would improve prediction of which items were subsequently recalled. This is the same approach adopted in previous eCMR simulations (Talmi, Lohnas & Daw, 2019), but here we report quantitative fits for the first time. Next, we asked whether scaling learning of each item with early LPP could improve, or indeed fully replace, that shared adjustment. Finally, we investigated whether LPP-bsed scaling was better applied to both emotional and neutral items or only to emotional items. We therefore crossed the presence or absence of preferential emotional learning, simulated via the increase in learning rate, with four LPP specifications: no LPP modulation, one modulatory relationship (shared across emotional and neutral items), a modulatory relationship limited to emotional items, or both overall and an emotion-specific modulatory relationships together. This crossing produced eight models. All eight models used the same memory architecture. We first describe that shared architecture and show where preferential emotional learning and LPP altered learning strength. We then describe model fitting, compare the eight models, and use simulations to determine which observed patterns the fitted models reproduced.

### eCMR Model Specification

We modeled recall with an adaptation of emotional Context Maintenance and Retrieval (eCMR) (Talmi et al., 2019). Like other retrieved-context models (Howard & Kahana, 2002; Polyn et al., 2009), eCMR represents context as an evolving internal state shaped by recently processed events. A studied item changes this state and forms associations in both directions: the item can later reinstate its study context, and the current context can later support retrieval of the item. Because a recalled item reinstates context before the next recall, stronger context-to-item learning can change which items enter the recalled set.

The context representation contained temporal and source components,

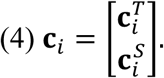

Temporal context represented the evolving sequence of events, whereas source context represented whether an item was emotional or neutral. These were components of one context representation rather than separate memory systems. All eight models represented both source categories. This means that models without preferential emotional learning therefore still distinguished emotional from neutral items.

Within each list, a feature vector **f***_i_* uniquely identified each studied item *i*. Two associative mappings connected item features and context. The feature-to-context mapping *M^FC^* allowed an item to reinstate context, whereas the context-to-feature mapping *M^CF^* allowed context to support candidate items. On each study trial, the item first generated temporal and source contextual inputs. For either component, C^IN^_i_ denotes the input generated by item *i*, and context updated as follows:

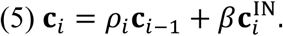

Here, *β* controlled how strongly the current item changed context, and *ρ_i_* kept context at unit length. Temporal and source context used separate encoding-drift parameters. Only after both components had been updated did the model learn the item’s associations, binding the item to a context state it had already helped create. Feature-to-context learning stored the updated context that the item could later reinstate. Context-to-feature learning determined how strongly that context would later support the item.

The temporal and source blocks of *M^CF^*, *M^CF^_T_* and *M^CF^_S_*, were updated as follows:

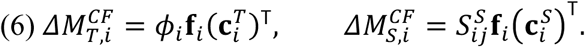

The primacy gradient *φ_i_* = 1 + *φ_s_*exp[−*φ_d_*(*i* − 1)] temporarily strengthened learning for early list positions. The focal emotional learning and LPP multipliers entered only the source learning strength *S^S^*, allowing them to be compared through the same pathway while temporal learning remained unchanged.

At recall onset, both context components shifted toward their start-of-list states. Temporal and source context then provided additive support *a_i_* for each candidate item:

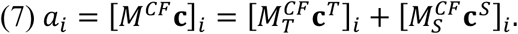

Among the studied items not yet recalled, choice sensitivity *τ_c_* converted this support into the probability of selecting item *i* next:

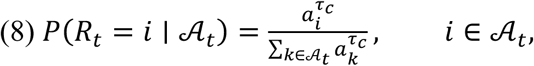

where A*_t_* contains the items still available at recall step *t*. After each selection, the item was removed from this set and reinstated its temporal and source context, changing the cue for the next recall. This cycle links source learning strength to an item’s probability of entering the recalled set.

The present model was a focused adaptation of published eCMR. Item-specific semantic associations were omitted because no validated similarity matrix was available for these stimuli. We also omitted source-switch disruption, which describes how transitions between emotional and neutral items alter temporal context, because this mechanism was outside the present comparison of learning-strength rules. Source drift was treated differently during study and recall because the two phases were differently constrained by the data. The observed study sequence allowed source-context drift during encoding to be estimated: smaller values allowed source information from preceding items to persist, whereas larger values made the current item more strongly determine source context. Recall order was not recorded, so the data did not directly reveal how source context carried over from one recall to the next. We therefore fixed source reinstatement during recall at 1, so each recalled item fully reinstated its learned source context. This setting can influence how one recall cues the next, but it was held constant across models while we compared how emotion and LPP altered source learning.

A separate choice concerned recall totals. The present comparison was designed to test whether alternative learning rate rules changed which items entered the recalled set. Predicting how many items were recalled would additionally require choosing a termination rule that converts the current state of retrieval into a decision to continue or stop. Different termination rules can map the same pattern of item strengths onto different recall totals, so including such a rules would combine the encoding question with a separate question about stopping. We therefore used the normalized selection rule above in place of the published accumulator and stopping process and conditioned the likelihood on each list’s observed recall count. For a list with *k* recalled items, the model evaluated which *k* items were recalled, not why recall ended after *k*. The comparison therefore concerns the composition of the recalled set rather than the mechanisms determining recall totals.

### Simulating encoding strength in eCMR

Following the attention-category variant of eCMR (Talmi et al., 2019), every model estimated a baseline strength for associations from source context to items, *ωφ_i_*. Here, *ω* scaled those associations relative to associations from temporal context to items, and *φ_i_* supplied the standard CMR primacy gradient (Sederberg et al., 2008; Polyn et al., 2009). Preferential emotional processing was assigned to source rather than temporal learning. If emotional processing instead strengthened temporal learning, emotional pure lists should show greater temporal clustering than neutral pure lists, a result for which there is currently no empirical support (Talmi et al., 2019).

The source learning rate *ωφ_i_* was further scaled by *φ*_emot_, a term we refer to as the “emotional multiplier”. This multiplier applied the same adjustment to every emotional item. Models without preferential emotional learning fixed the emotional multiplier *φ*_emot_ at 1, so the multiplier made no adjustment. Models with preferential emotional learning estimated *φ*_emot_ within the range [0, 10]. Values above 1 strengthened source learning for emotional items relative to neutral items, whereas values below 1 weakened it. The upper bound was a broad numerical constraint rather than a theoretical ceiling: it permitted as much as a tenfold increase, while the estimates in the models reported here ranged from 1.86 to 2.66. Because the multiplier applied the same adjustment to every emotional item, it does not represent variation in learning rate among them.

To test whether LPP could explain learning rate variations among encoded items, we used the standardized LPP elicited by each study item at the early time window (henceforth: seLPP for short) to scale the same source learning term. We denote the seLPP for item i in list j by zᵢⱼ and coded item category as *ei* = 1 for emotional items and *ei* = 0 for neutral items. Source learning strength therefore combined the ordinary source-learning baseline, the emotional-learning multiplier, and the seLPP term:

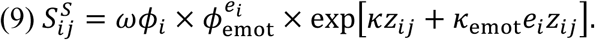

Within the seLPP term, *κ* described how seLPP changed source learning for both emotional and neutral items. For emotional items, *κ*_emot_ was added to *κ* . Neutral items therefore received the multiplier exp(*κz_i_*_j_), whereas emotional items received exp\*κz_i_*_j_ + *κ*_emot_*e_i_z_i_*_j_’.

The exponential form kept the learning rate positive and expressed the LPP effect as a proportional change. When *z_i_*_j_ = 0, the seLPP term equaled 1 and did not alter learning rates.

We evaluated four LPP specifications. “No LPP” fixed both *κ* and *κ*_emot_ at zero. “General LPP” estimated *κ* alone, allowing seLPP to modify source learning in both emotional and neutral items through one shared coefficient. “Emotion-dependent LPP” estimated *κ*_emot_ alone, allowing seLPP to modify source learning only for emotional items. These two specifications each added one parameter and were equally complex. “Combined LPP” estimated both coefficients, combining an seLPP coefficient shared by both categories with an additional coefficient for emotional items. The Combined LPP specification most closely mirrored the empirical binomial generalised mixed-effect model (equation 2), which included both a general seLPP coefficient and an interaction between emotion and seLPP, although the follow-up analyses found a significant seLPP–memory relationship for emotional but not neutral items.

### Model comparison

To compare the proposed learning mechanisms, we combined two settings of the emotional multiplier with the four seLPP specifications described above. In one setting, ϕₑₘₒₜ was fixed at 1, so there was no common adjustment to source learning for emotional items. Source learning could nevertheless vary between items through seLPP where the specification allowed it. In the second setting, ϕₑₘₒₜ was estimated, allowing every emotional item to receive the same adjustment in addition to any modulation by seLPP. Crossing these two settings with the four seLPP specifications produced the eight models in Table 1.

**Table 1:** Pooled fit of the eight eCMR model variants formed by crossing seLPP modulation with a fixed or estimated emotional learning multiplier.

| Model number | Preferential emotional learning | seLPP specification | Number of Free parameters | NLL | AIC | ΔAIC |
| --- | --- | --- | --- | --- | --- | --- |
| 1 | No | None | 11 | 8312.40 | 16646.80 | 133.904 |
| 2 | No | General | 12 | 8300.54 | 16625.08 | 112.178 |
| 3 | No | Emotion-dependent | 12 | 8275.47 | 16574.94 | 62.035 |
| 4 | No | Combined LPP<sup>a</sup> | 13 | 8275.47 | 16576.94 | 64.035 |
| 5 | Yes | None | 12 | 8258.46 | 16540.92 | 28.021 |
| 6 | Yes | General | 13 | 8246.00 | 16518.00 | 5.096 |
| 7 | **Yes** | **Emotion-dependent** | **13** | **8243.45** | **16512.90** | **0.000** |
| 8 | Yes | Combined LPP<sup>a</sup> | 14 | 8243.45 | 16514.90 | 2.000 |

Note: All models contained temporal context and separate source representations for emotional and neutral items. Estimating the multiplier tested the directional hypothesis of preferential emotional learning while also permitting attenuation. In the Preferential emotional learning column, “No” indicates that the emotional learning multiplier was fixed at 1; Yes indicates that it was estimated. p is the number of free parameters; NLL is the retained negative log-likelihood; ΔAIC is relative to the lowest AIC. Bold identifies the lowest-AIC model. <sup>a</sup> For both Combined LPP models, the direct searches did not improve upon the corresponding Emotion-dependent solution, so that nested solution was retained; the number of free parameters and AIC nevertheless include the Combined LPP model’s additional parameter.

Table 1 can be read in two ways. Within each ‘preferential learning’ group, i.e. when preferential learning is fixed to “no” or “yes”, the LPP specification changes while the treatment of the emotional learning multiplier stays the same. Across preferential learning groups, models with the same LPP specification differ only in whether the emotional learning multiplier is fixed or estimated; this comparison tests preferential emotional learning. The No LPP model provides the baseline for testing whether seLPP improves prediction. Comparing General LPP with Combined LPP tests whether emotional items have an additional LPP effect beyond the relationship shared by both categories. Comparing Emotion-dependent LPP with Combined LPP tests whether the shared relationship is also needed.

Table 2 summarizes the parameters and the constraints used to define the eight models. Eleven parameters were estimated in every model; source context reinstatement during recall was fixed at 1. Depending on the model, the emotional learning multiplier and the two seLPP coefficients were either fixed at their neutral values or estimated. The models therefore contained between 11 and 14 free parameters, with General LPP and Emotion-dependent LPP equally complex within each preferential-learning setting (12 parameters without preferential learning and 13 with it).

**Table 2:** Parameters and constraints used in the eight-model specification.

| Parameter | Role | Fitted range or fixed value |
| --- | --- | --- |
| $\beta_{\text{enc}}^T$ | Temporal-context drift during study | (0,1) |
| $\beta_{\text{enc}}^S$ | Source-context drift during study | (0,1) |
| $\beta_{\text{start}}$ | Drift toward start-of-list context before recall | (0,1) |
| $\beta_{\text{rec}}^T$ | Temporal-context reinstatement after recall | (0,1) |
| $\beta_{\text{rec}}^S$ | Source-context reinstatement after recall | 1 (fixed) |
| $\alpha$ | Pre-existing support from context shared across items | (0,100] |
| $\delta$ | Pre-existing support from context specific to each item | (0,100] |
| $\gamma$ | Rate of learning associations from items to context | (0,1) |
| $\phi_s$ | Primacy scale | (0,100] |
| $\phi_d$ | Primacy decay | (0,100] |
| $\tau_c$ | Sensitivity of recall choices to activation | (0,100] |
| $\omega$ | Source learning relative to temporal learning | (0,10] |
| $\phi_{\text{emot}}$ | Multiplier on source learning for emotional items | 1 (fixed) or (0,10] |
| $\kappa$ | seLPP coefficient shared by emotional and neutral items | 0 (fixed) or [0,0.1923] |
| $\kappa_{\text{emot}}$ | Additional seLPP coefficient for emotional items | 0 (fixed) or [0,0.1923] |

The seLPP coefficients *κ* and *κ*_emot_ were constrained to [0, 0.1923]. The lower bound expressed the directional prediction that larger seLPP would not weaken learning. The upper bound was chosen so that, at the largest seLPP value in the modeling data (11.9711), a single coefficient could increase learning strength by at most a factor of 10: exp(0.1923 × 11.9711) = 10. Having defined the alternatives, we next describe the common evidence used to compare them.

### Pooled estimation and evaluation

The modeling dataset comprised 6,840 studied items from 342 lists contributed by 38 participants, with recall status recorded for every item. Usable seLPP measurements were available for 6,469 items. For the remaining 371 items (5.4%), we imputed seLPP without using recall status. Missing values were replaced, in order, by the mean for the same emotional category on that participant’s list, the mean for the complete list, or the corresponding category mean across the dataset. We fitted one parameter vector to all 342 lists (each participant contributed nine lists) such that the resulting estimates describe the pooled pattern across participants.

The records identified each list’s recalled items but not their output order. Let *n*_j_ = gS_j_g be the number of items in recalled set S_j_, let *π*_j*k*_ be its *k*th sampled order, and let **θ** be the model parameters. Because reinstatement makes eCMR sequence-dependent, the probability of a recalled set is the sum of the probabilities of its *n*_j_! possible orders. We approximated this sum using *K* = 50 fixed orders sampled uniformly:

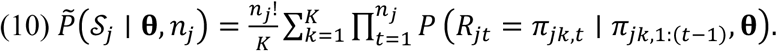

Each product is the probability of one possible recall sequence; multiplying their sampled average by *n*_j_! estimates the sum over all possible recall orders. For fitting, we omitted *n*_j_! because it depends only on the observed set size. This omission increases the NLL for list *j* by logN*n*_j_!O, a parameter-independent constant shared by every model; parameter estimates, NLL differences, AIC differences, and model rankings are therefore unchanged. The reported NLL and AIC values are based on this constant-omitted objective. The sampled orders were generated with seed 0 and held fixed for every model. Because the likelihood also excluded termination, it predicts recalled-item identity conditional on the observed number of recalls rather than explaining how many items were recalled.

We minimized summed NLL with differential evolution (Storn & Price, 1997), using 15 candidate vectors per free parameter for at most 1,000 generations. Three independent starts were run for each model, and the lowest NLL was retained. If the best search result for a larger model had a higher NLL than any immediately nested simpler model, we followed Morton and Polyn (2016) by evaluating those simpler-model solutions in the larger parameterization, with the added parameter set to its neutral value, and retained the candidate with the lowest NLL. We compared best-found solutions using AIC = 2NLL + 2*p*, where *p* is the number of free parameters (Burnham & Anderson, 2002).

To interpret the differences between fits, we generated 200 complete datasets from each fitted model using the observed lists, standardized seLPP values, and recall counts. We summarized the difference in recall between emotional and neutral items and the difference in seLPP between remembered and forgotten items within each category. Events whose seLPP values were imputed were excluded from the latter diagnostic. These summaries were consequences of which items the model recalled, not additional fitting targets, and therefore provide explanatory diagnostics rather than out-of-sample validation.

## Results

### Overall comparison

Table 1 compares how well the eight source-context models predicted which items entered each list’s recalled set, given the observed number of recalls. NLL measures prediction error, so lower values indicate better prediction. AIC also penalizes models for additional free parameters, and ΔAIC is the difference between each model’s AIC and the lowest values.

The model combining preferential emotional learning with an seLPP modulation only for emotional items had the lowest AIC (NLL = 8243.450, *p* = 13, AIC = 16512.900). The Combined LPP model, which added a general seLPP coefficient, retained the same NLL but used one additional parameter and therefore had an AIC two points higher.

Figures 5 and 6 show whether the fitted mechanisms reproduced the two empirical patterns: the recall advantage for emotional items and the stronger relationship between seLPP and memory for emotional than neutral items. We compare the observed data with simulations from Emotion-dependent LPP (model 3), Preferential Learning (model 5), and their combination (Preferential learning + Emotion-dependent LPP, model 7). Figure 5 shows whether each model produces the observed recall advantage for emotional over neutral items. Figure 6 shows whether each model preferentially recalls items with larger seLPP values and whether this relationship differs between emotional and neutral items. These summaries were calculated after fitting and did not contribute to NLL or AIC.

**Figure 5.**
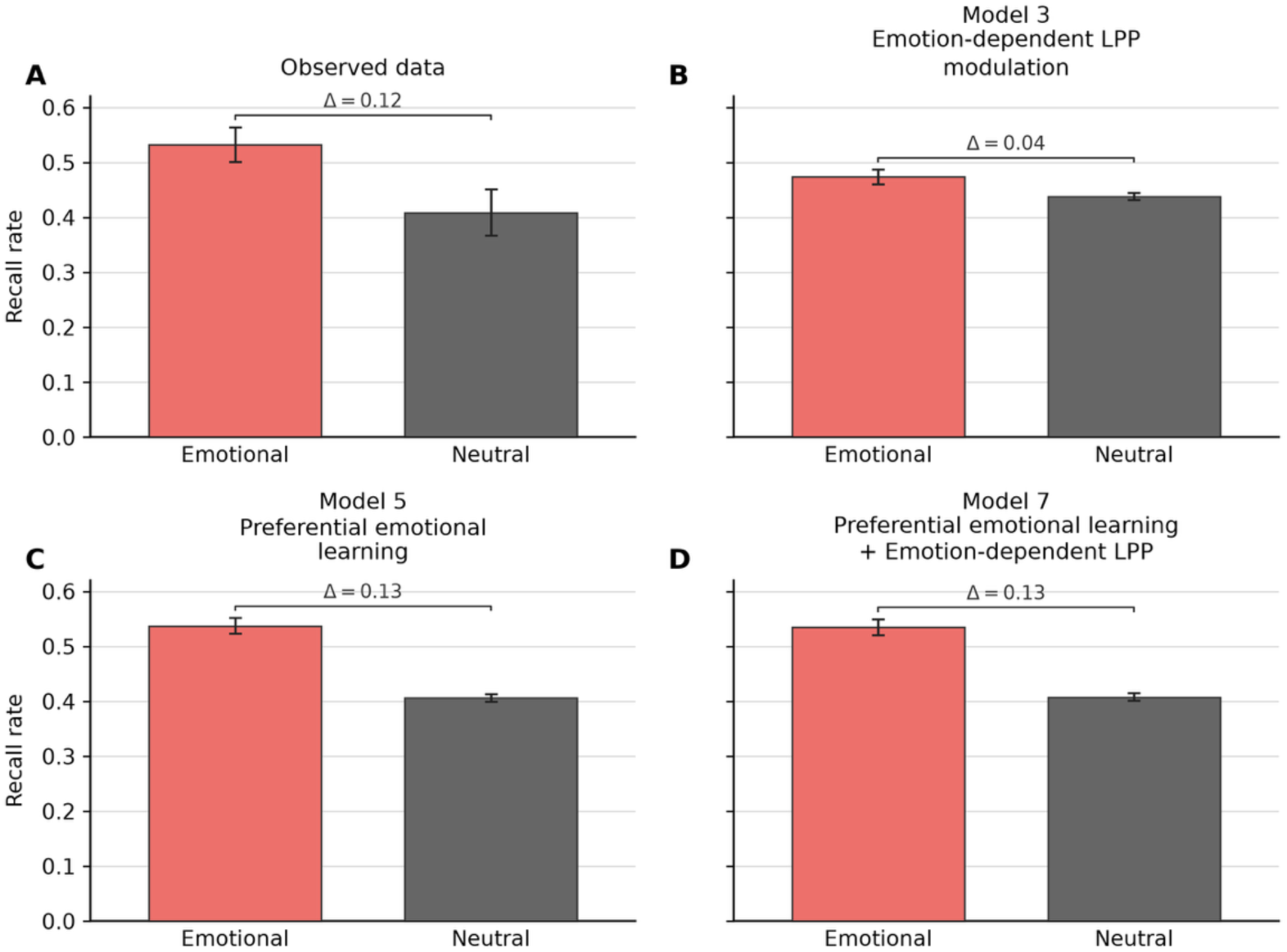
Observed and model-predicted recall rates for emotional and neutral items. Panels show (A) observed data, (B) Emotion-dependent LPP alone (model 3), (C) preferential emotional learning alone (model 5), and (D) both mechanisms together (model 7). Brackets show the difference between emotional and neutral items. Observed error bars are 95% participant-cluster bootstrap confidence intervals; predicted error bars are central 95% intervals across 200 complete simulated datasets.

**Figure 6.**
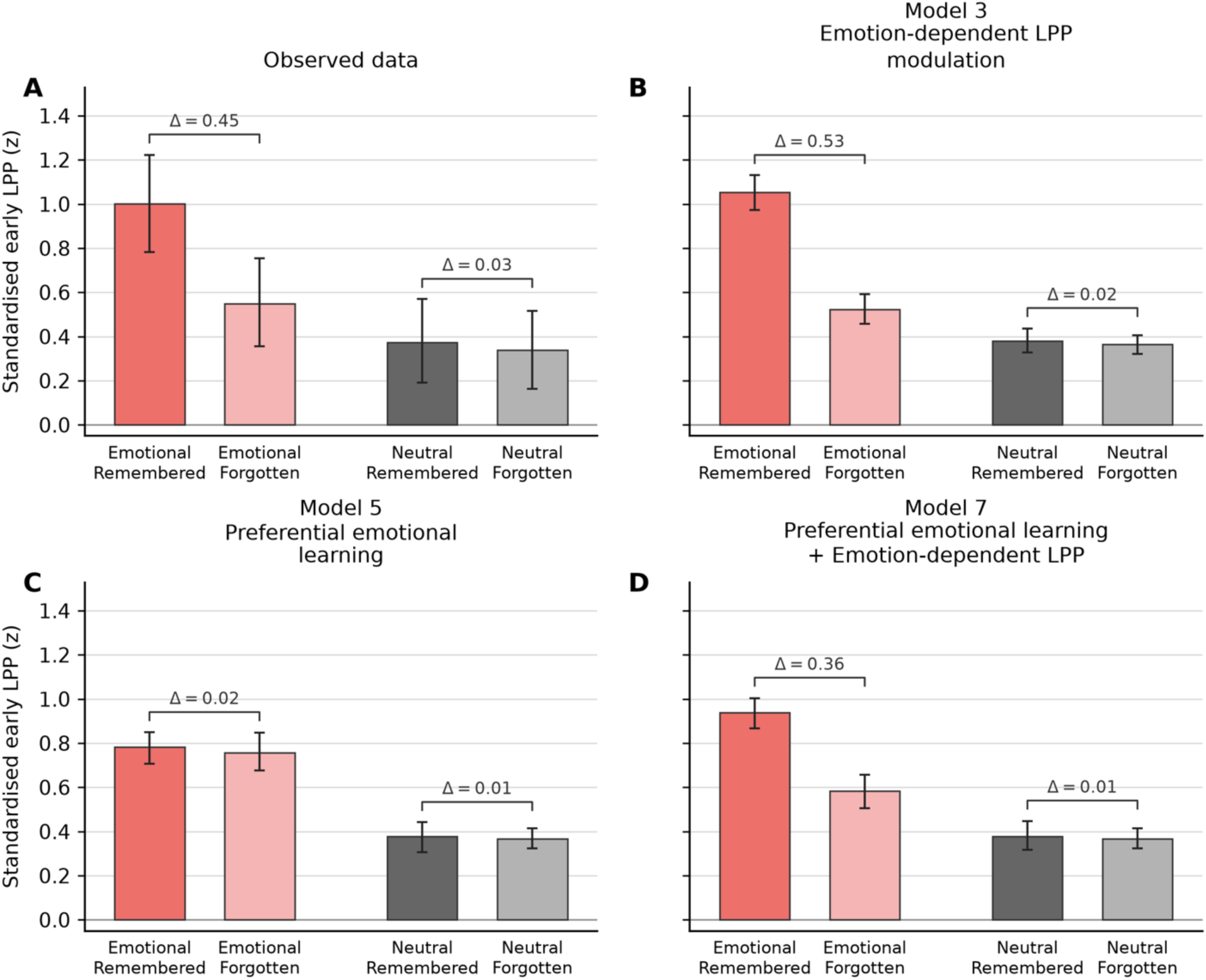
Observed and model-predicted standardized seLPP by item category and memory status. Panels show (A) observed data, (B) Emotion-dependent LPP alone (model 3), (C) preferential emotional learning alone (model 5), and (D) both mechanisms together (model 7). *Brackets show the difference between remembered and forgotten items within each category. Observed error bars are 95% participant-bootstrap confidence intervals; predicted error bars are central 95% intervals across 200 complete simulated datasets.* Amplitudes were standardised separately for each trial and electrode using the mean and standard deviation of the −200 to 0 ms prestimulus baseline.

First, we asked whether preferential emotional learning improved prediction of which items were recalled. In these models, preferential emotional learning allowed all emotional items to share an increase in source learning, in addition to any modulation by LPP amplitude during their encoding. Estimating the emotional learning multiplier rather than fixing it at 1 lowered AIC under every seLPP specification, by between 62.035 and 107.082 AIC units. The estimated multiplier exceeded 1 in all four models (range = 1.86–2.66). Figure 5 shows the corresponding simulated recall patterns. The observed difference in recall rate between emotional and neutral items was 12.4 percentage points. The model with Emotion-dependent LPP modulation but no preferential emotional learning produced a difference of only 3.6 percentage points, whereas preferential emotional learning alone and preferential emotional learning combined with Emotion-dependent LPP modulation produced differences of 13.1 and 12.7 percentage points, respectively. Thus, trial-level LPP differences alone did not reproduce the overall recall advantage for emotional items. Allowing emotional items to increase learning improved prediction even when LPP modulation was available.

Second, we asked whether seLPP improved the model’s prediction of which items entered the recalled set, given the observed number of recalled items. Each of the three LPP specifications lowered AIC relative to the corresponding model without LPP modulation, suggesting that adding the neural evidence into this mechanistic model helps predict memory recall. The improvements ranged from 21.727 to 71.869 AIC units when preferential emotional learning was absent and from 22.926 to 28.021 AIC units when it was present. Figure 6 illustrates how this improvement was realized. In the observed data, mean seLPP was 0.455 z-score units higher for remembered than forgotten emotional items, but 0.035 units higher for remembered than forgotten neutral items. Preferential emotional learning alone produced almost no difference by memory status for either item category (0.024 for emotional and 0.010 for neutral), because its learning multiplier did not vary among items within a category. With Emotion-dependent LPP modulation, seLPP was substantially higher for remembered than forgotten emotional items but differed little by memory status for neutral items, both without preferential emotional learning (0.530 and 0.016) and with it (0.356 and 0.011). Each simulated item retained its observed seLPP value, so this pattern occurred because LPP modulation made emotional items with larger values more likely to enter the recalled set. seLPP therefore contributed information about differences among emotional items that the common emotional learning multiplier did not provide.

Finally, we compared the alternative ways in which seLPP could affect learning. Emotion-dependent LPP had a lower AIC than General LPP by 50.143 points without preferential emotional learning (compare models 3 to 2) and by 5.096 points with it (compare models 7 to 6), suggesting that Emotion-dependent LPP was the better model regardless of the presence of preferential emotional learning. Adding the general coefficient to the Emotion-dependent specification produced no further improvement in the retained NLL; the Combined LPP models (4 and 8) therefore had AIC values exactly two points higher that Emotion-dependent models (3 and 7) because they contained one additional parameter. Matching the empirical results, where the main effect of seLPP was qualified by a significant interaction with the Emotion factor and did not significantly predict recall in an analysis focused on the neutral items, the available optimization did not find evidence that a general seLPP coefficient improved likelihood once the coefficient applied only to emotional items was included. These results indicate that trial-level seLPP was particularly informative for predicting which emotional items were recalled.

Among the equally complex 12-parameter models, preferential emotional learning alone also had a lower AIC than General LPP alone by 84.156 points and Emotion-dependent LPP alone by 34.014 points. seLPP therefore added information about which items were recalled but did not replace preferential emotional learning as an account of the overall recall advantage for emotional items.

## Discussion

The model comparison indicated that preferential emotional learning and seLPP provided complementary information. Preferential emotional learning accounted for the overall recall advantage for emotional items, whereas the seLPP improved prediction of which individual items entered the recalled set given each list’s observed recall count. This contribution was concentrated among emotional items: the Emotion-dependent specification outperformed the equally complex General specification, and the model with the lowest AIC combined Emotion-dependent LPP modulation with preferential emotional learning. Trial-level seLPP therefore refined predictions within the emotional condition rather than replacing the emotion-dependent mechanism proposed by eCMR.

The Emotion-dependent + preferential learning model specification (model 7) was consistent with the emotion-specific seLPP–memory relationship found in the empirical analyses: it included one coefficient applied to all emotional items and one seLPP coefficient that differentiated between them as a function of early LPP during encoding. The available optimization did not find a likelihood improvement from adding the shared seLPP coefficient once a coefficient applied only to emotional items was included. Because the Combined LPP model contained one additional parameter, the Emotion-dependent model had the lower AIC.

## General Discussion

This project investigated whether the LPP, measured during encoding, is suitable for neuro-cognitive modelling of emotional memory recall. The ERP analyses showed that the early LPP was reliable in a higher proportion of the participants than the late LPP and was related to subsequent memory. The early LPP predicted memory in an emotion-specific and level-dependent way, tracking recall only for emotional items and only at the single-trial level, which means that the LPP captures important variations between the effect of each picture on subsequent memory. The modelling results showed that trial-level early LPP complemented eCMR’s common learning-rate increase for all emotional items, with the best-fitting model including both. Taken together, the results show that single-trial LPP data can help predict memory performance and support the conceptualisation of learning proposed in eCMR.

At the group-level, the focal and the more conservative global ERP analyses replicated the typical LPP ERP effect and Zarubin and colleague’s (2020) original report with more positive amplitudes for emotional compared to neutral stimuli over centro-parietal electrode sites. The individual-level reliability was higher for the early LPP (73.7% of the participants) compared to the late LPP (50% of the participants). The reliability of the early LPP here was lower than what has been observed in a previous experiment (Schupp & Kirmse 2021). This confirms our initial concern that intentional encoding could dilute the emotional modulation of this signal, for example, if participants redistribute attention while comparing or organising items for later recall. Another possible reason for the lower reliability of the LPP in the present study is the stimuli. A recent study showed that while the LPP was reliable for an erotic picture in 93.8% of their sample, the LPP to a mutilation image, which were more similar to the emotionally-valenced images in this study, was only seen in 62.5% of the participants, between the reliability values for the early and late LPP in our study (Schupp, Flösch & Flaisch, 2025). Other factors that may cause variability in the reliability of the LPP across studies are presentation duration and the signal-to-noise ratio (e.g. number of trials) and the number of participants included in the analysis. Pleasingly, however, despite these challenges, the results of the reliability analysis suggest that the early LPP is reliable in a standard emotional memory encoding task, with typical presentation timing and stimuli, and that it is reliable in more participants than the late LPP.

Besides being reliable in a higher proportion of the participants, the early LPP also showed a relation to subsequent memory, in the focal ERP analysis (Fig. 2 and 4E), which was absent in the late LPP, replicating previous studies linking the early LPP to subsequent memory (Barnacle et al., 2018; Dolcos & Cabeza, 2002; Watts et al., 2014; Zarubin et al., 2020). The global ERP analysis (Fig. 3) showed that there was an Emotion × Memory interaction in a time window roughly overlapping with the early LPP time window (360-1240 ms compared to the 400-1000 ms used for the focal early LPP analysis). The early LPP has spatiotemporal overlap with the P300 component and has been suggested to reflect the same underlying neurocognitive mechanisms as the P300 (for reviews see Fields 2023; Hajcak & Foti 2020).

Interestingly, both the global ERP analysis and the focal analysis of the early LPP time window showed that LPP amplitude only predicted subsequent memory for emotional pictures. The model comparison showed the same specificity: the Emotion-dependent LPP specification had a lower AIC than the equally complex General specification. This convergent pattern appears to be inconsistent with other theoretical accounts of the LPP. Some suggest that the LPP reflects attention, stimulus significance (Hajcak & Foti, 2020) or tagging of memories as important for consolidation (Fields, 2023). It is difficult to reconcile the pattern of results we observed here with these theories, since they predict a subsequent memory LPP effect also for neutral items, not just for emotional ones. We speculate that potential differences in the significance of neutral stimuli, which would normally capture attention and promote subsequent memory, were overshadowed by the presence of highly emotional stimuli. Future research can test this hypothesis by examining lists that consist only of stimuli of one type (pure lists) compared to those that mix emotional and neutral stimuli (mixed lists). We predict that in pure lists, the LPP is larger for subsequently remembered items in both emotional and neutral lists.

We next investigated the relationship between the LPP and emotion-enhanced memory. We found moderate Bayesian support for no correlation between the LPP (negative-neutral) and emotion-enhanced memory between participants in either the early or the late LPP time window (Fig. 4B and 4C). Individual differences in the magnitude of the LPP could be dependent on several factors unrelated to the cognitive processes the LPP reflects. Among others, differences in neuroanatomy, skull thickness and electrode impedance could affect LPP amplitude and mask an underlying relationship. Similarly, individual differences in amplitude of the left parietal old/new effect in recognition memory have been shown not to correlate with individual differences in recollection despite showing a relationship within participants (MacLeod & Donaldson, 2017). More surprisingly, we also observed moderate Bayesian evidence for no correlation between the amplitude of the LPP and the size of the emotion-enhanced memory effect across the nine lists within participants. Instead, the relationship was present when analysing the data at the single-trial level using a binomial generalized mixed-effects model.

Taken together, the pattern of results suggests that variation between processing of individual stimuli are particularly influential for their eventual recall. The finding that the relationship between the emotion-enhanced LPP and emotion-enhanced memory was best captured at the single-trial level informed our use of trial-level early LPP to simulate which encoded items were later recalled.

Computational models are useful because they specify how encoding, maintenance, and retrieval processes may give rise to recall. Comparing their predictions with observed recall allows alternative computational accounts to be evaluated against one another. Here, the best-fitting model allowed a neural signal measured during encoding – the early LPP – to modulate source learning specifically for emotional items, alongside the common learning-rate increase for all emotional items. This finding suggests that the LPP may reflect item-level variation in encoding processes that eCMR represents as stronger associations between emotional source context and individual items, rather than merely reflecting a general emotional response.

More broadly, these results provide a proof of principle that trial-level early LPP can add item-level information to a computational account of emotional recall. This improvement concerned which emotional items entered the recalled set, conditional on each list’s observed recall count. It does not yet establish prospective prediction for individuals, since the model used one pooled parameter vector, did not model recall termination, and was evaluated on the same data used for fitting rather than on held-out data. Individualized prediction would require participant-level or hierarchical modelling, a validated account of recall termination, and evaluation on new participants or lists. With those extensions, and building on present results, scalp EEG and eCMR could be used jointly to predict which items will come to mind for an individual participant even without requiring explicit verbal recall. Such ability would be invaluable in a range of practical situations including clinical, educational, and commercial settings. For example, if a language tutor knew which words their pupil was most likely to recall later, they could prioritise the revision of the more challenging words. Crucially, the benefit is not only practical. Requiring the parameter (Φ_emot) to answer to an independent neural measure yields a more principled account than free estimation (Turner et al., 2016), and here it yielded a specific one: the LPP predicted recall for emotional items alone and improved the model alongside preferential emotional learning rather than instead of it.

## Author contributions

**Robin Hellerstedt**: Conceptualisation, data curation, methodology, formal analysis, project administration, visualisation, writing – original draft, writing – review & editing. **Jordan Gunn**: Conceptualisation, formal analysis, methodology, software, visualisation, writing – original draft, writing – review & editing. **Deborah Talmi**: Conceptualisation, funding acquisition, methodology, project administration, resources, supervision, visualisation, writing – original draft, writing – review & editing.

## Supplementary material

**Table 4:**
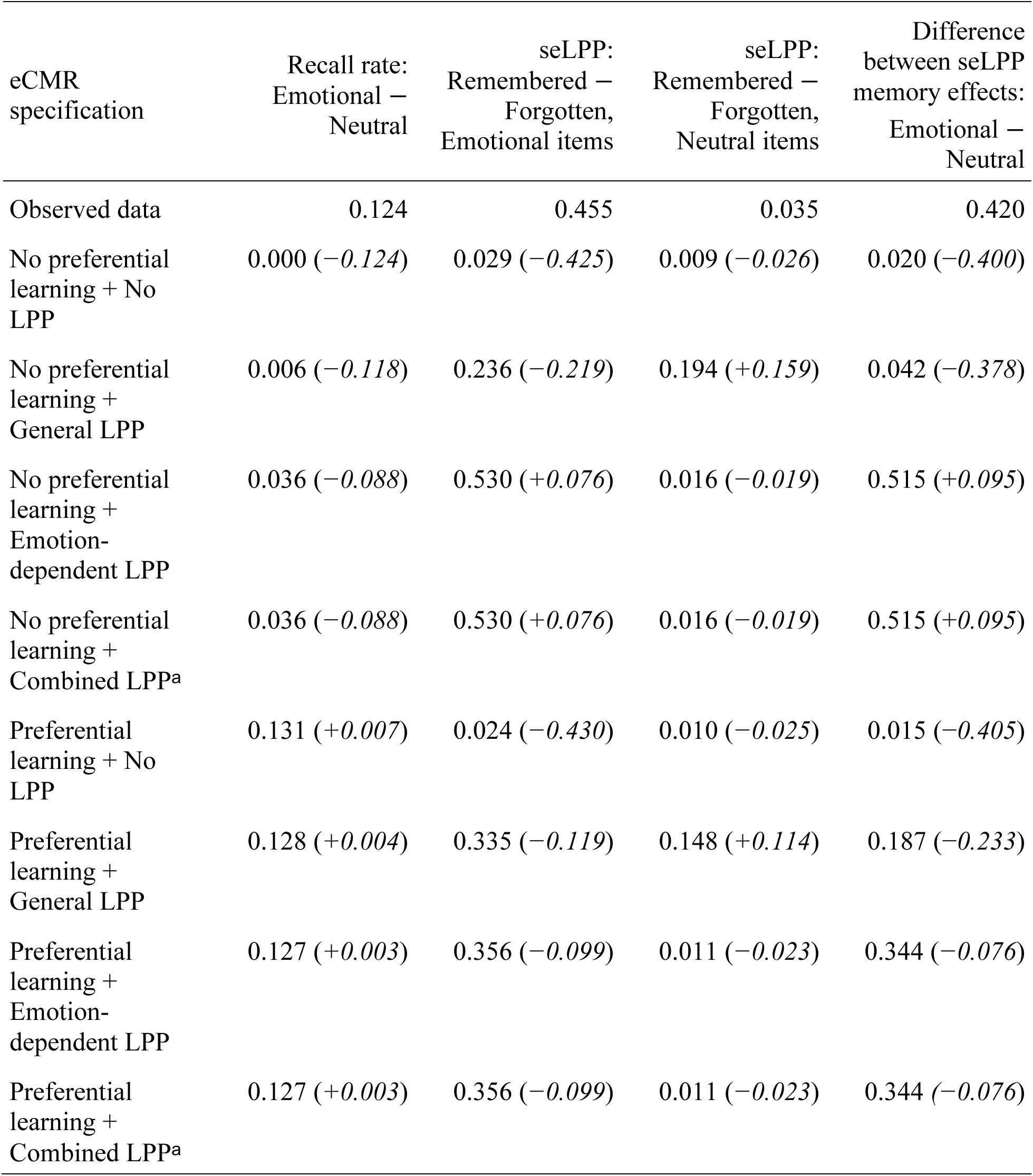
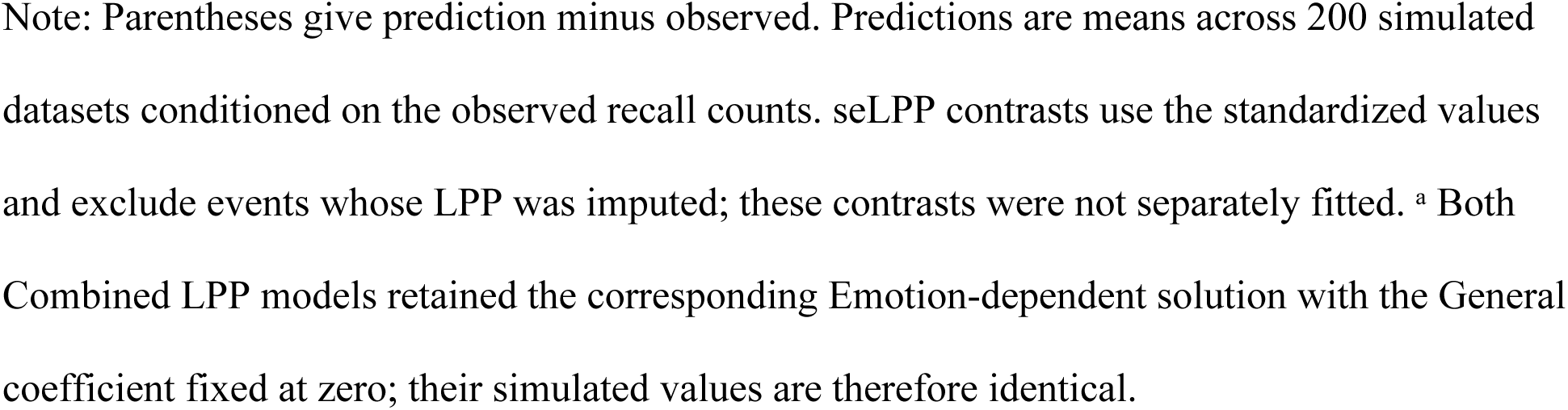
Observed and model-predicted contrasts in recall rate and standardized seLPP.

| eCMR specification | Recall rate:<br>Emotional –<br>Neutral | seLPP:<br>Remembered –<br>Forgotten,<br>Emotional items | seLPP:<br>Remembered –<br>Forgotten,<br>Neutral items | Difference<br>between seLPP<br>memory effects:<br>Emotional –<br>Neutral |
| --- | --- | --- | --- | --- |
| Observed data | 0.124 | 0.455 | 0.035 | 0.420 |
| No preferential<br>learning + No<br>LPP | 0.000 (–0.124) | 0.029 (–0.425) | 0.009 (–0.026) | 0.020 (–0.400) |
| No preferential<br>learning +<br>General LPP | 0.006 (–0.118) | 0.236 (–0.219) | 0.194 (+0.159) | 0.042 (–0.378) |
| No preferential<br>learning +<br>Emotion-dependent LPP | 0.036 (–0.088) | 0.530 (+0.076) | 0.016 (–0.019) | 0.515 (+0.095) |
| No preferential<br>learning +<br>Combined LPP<sup>a</sup> | 0.036 (–0.088) | 0.530 (+0.076) | 0.016 (–0.019) | 0.515 (+0.095) |
| Preferential<br>learning + No<br>LPP | 0.131 (+0.007) | 0.024 (–0.430) | 0.010 (–0.025) | 0.015 (–0.405) |
| Preferential<br>learning +<br>General LPP | 0.128 (+0.004) | 0.335 (–0.119) | 0.148 (+0.114) | 0.187 (–0.233) |
| Preferential<br>learning +<br>Emotion-dependent LPP | 0.127 (+0.003) | 0.356 (–0.099) | 0.011 (–0.023) | 0.344 (–0.076) |
| Preferential<br>learning +<br>Combined LPP<sup>a</sup> | 0.127 (+0.003) | 0.356 (–0.099) | 0.011 (–0.023) | 0.344 (–0.076) |

Note: Parentheses give prediction minus observed. Predictions are means across 200 simulated datasets conditioned on the observed recall counts. seLPP contrasts use the standardized values and exclude events whose LPP was imputed; these contrasts were not separately fitted. <sup>a</sup> Both Combined LPP models retained the corresponding Emotion-dependent solution with the General coefficient fixed at zero; their simulated values are therefore identical.

**Table 5.** Retained pooled parameter estimates for the eight source-context models.

**Panel A. Models without preferential emotional learning**
| Parameter | No LPP | General LPP | Emotion-dependent LPP | Combined LPP<sup>a</sup> |
| --- | --- | --- | --- | --- |
| $\beta_{\text{enc}}^T$ | 0.71851 | 0.78142 | 0.80691 | 0.80691 |
| $\beta_{\text{enc}}^S$ | 0.060535 | 0.024017 | 0.036409 | 0.036409 |
| $\beta_{\text{start}}$ | 0.99726 | 0.99975 | 0.99975 | 0.99975 |
| $\beta_{\text{rec}}^T$ | 0.42503 | 0.49575 | 0.33069 | 0.33069 |
| $\beta_{\text{rec}}^S$ | [1] | [1] | [1] | [1] |
| $\alpha$ | 0.30498 | 1.5303 | 0.72775 | 0.72775 |
| $\delta$ | 23.840 | 53.847 | 7.4796 | 7.4796 |
| $\gamma$ | 0.28870 | 0.24546 | 0.34176 | 0.34176 |
| $\phi_s$ | 1.3389 | 0.67787 | 0.28273 | 0.28273 |
| $\phi_d$ | 82.540 | 1.0314 | 1.4471 | 1.4471 |
| $\tau_c$ | 0.64638 | 1.0003 | 1.5895 | 1.5895 |
| $\omega$ | 1.8797 | 7.2810 | 1.5667 | 1.5667 |
| $\phi_{\text{emot}}$ | [1] | [1] | [1] | [1] |
| $\kappa$ | [0] | 0.10149 | [0] | 0 |
| $\kappa_{\text{emot}}$ | [0] | [0] | 0.16688 | 0.16688 |

**Panel B. Models with preferential emotional learning**
| Parameter | No LPP | General LPP | Emotion-dependent LPP | Combined LPP<sup>a</sup> |
| --- | --- | --- | --- | --- |
| $\beta_{\text{enc}}^T$ | 0.78546 | 0.86821 | 0.84297 | 0.84297 |
| $\beta_{\text{enc}}^S$ | 0.037635 | 0.058007 | 0.076622 | 0.076622 |
| $\beta_{\text{start}}$ | 0.99997 | 0.99993 | 0.99997 | 0.99997 |
| $\beta_{\text{rec}}^T$ | 0.14929 | 0.32673 | 0.19575 | 0.19575 |
| $\beta_{\text{rec}}^S$ | [1] | [1] | [1] | [1] |
| $\alpha$ | 7.7623 | 2.4052 | 2.8967 | 2.8967 |
| $\delta$ | 94.112 | 38.867 | 30.717 | 30.717 |
| $\gamma$ | 0.19099 | 0.21932 | 0.24409 | 0.24409 |
| $\phi_s$ | 0.43136 | 1.0300 | 0.51366 | 0.51366 |
| $\phi_d$ | 1.6925 | 1.1998 | 1.8352 | 1.8352 |
| $\tau_c$ | 1.8059 | 1.3229 | 1.6787 | 1.6787 |
| $\omega$ | 1.7292 | 1.4715 | 1.3416 | 1.3416 |
| $\phi_{\text{emot}}$ | 2.6595 | 2.4435 | 1.8613 | 1.8613 |
| $\kappa$ | [0] | 0.12845 | [0] | 0 |
| $\kappa_{\text{emot}}$ | [0] | [0] | 0.11006 | 0.11006 |

*Note.* Estimates are rounded to five significant digits; square brackets identify parameters fixed by the model specification. Parameter roles and fitted ranges are reported in Table 2. The Combined LPP models retained the corresponding Emotion-dependent solutions evaluated with $\kappa = 0$ ; all other estimates therefore repeat those columns. The unbracketed $\kappa = 0$ was estimated, not fixed.

